# Route Retracing: Way-pointing in Trail Following Ants

**DOI:** 10.1101/2023.09.02.556024

**Authors:** Cody A Freas, Marcia L Spetch

## Abstract

Maintaining positional estimates of goal locations is a fundamental task for navigating animals. Diverse animal groups, including both vertebrates and invertebrates, can accomplish this through path integration (PI). During PI, navigators integrate movement changes, tracking both distance and direction, to generate a spatial estimate of their start location, or global vector, allowing efficient direct return travel without retracing the outbound route. In ants, PI is accomplished through the coupling of pedometer and celestial compass estimates. Within the PI system, it has been theorized navigators may segment the global vector into local-vectors for way-pointing. However, this is controversial, as these navigators may instead be homing via the view alignment. Here, we present evidence trail-following ants can attend to segments of their global vector to retrace their non-straight pheromone trails, without the confound of familiar views. *Veromessor pergandei* foragers navigate via directionally distinct segments of their PI by orienting along separate legs of their inbound route at unfamiliar locations, indicating these changes are not triggered by familiar external cues, but by the PI state. These findings contrast with the view of path integration as a singular memory estimate and underscore the system’s ability to way-point to intermediate goals along the inbound route. We discuss how the foraging ecology of ant species that rely on non-straight pheromone-marked trails may support attending vector segments to remain on the pheromone rather than attempting straight-line shortcuts back to the nest.

## Introduction

Efficient homing to goals is a critical recurring challenge for navigating animals, often necessitating the navigator track its position in relation to these sites, even in habitats that are visually featureless. Insect navigators possess several strategies to efficiently orient to compass headings and reach goals (Heinze et al. 2018; Wehner 2020; Freas and Cheng 2022). One widely documented solution is path integration (PI), where the navigator maintains an updating positional estimate relative to its origin by measuring distance and directional movement along the route, integrating these into a global vector (Collett and Collett 2000; Wehner and Srinivasan 2003).

Many invertebrate navigators (ants, honeybees, bumblebees, flies, cockroaches, field crickets and crabs, spiders, etc.) are well documented to attend to a PI-derived vector to return directly, along a straight-line route to their nest/burrow rather than re-tracing their winding outbound paths (Cheng 2006; Collett and Collett 2000; Freas and Spetch 2023; Srinivasan 2011; Wehner and Srinivasan 2003; Zeil and Layne 2002). In ants, PI is modelled as a single estimate, integrating pedometer and celestial compass information to produce a global vector that directs returning individuals back to the nest, with a distinct lack of the ability to home to intermediate sites enroute. In visually cluttered environments, as individuals become experienced with the landmarks along their established foraging route, they develop stereotypical paths detouring around any obstacles (i.e., bushes or rocks). These detours often involve periods of turning away from the global vector before reorienting back to the nest direction (Collett et al. 1992; Bisch-Knaden and Wehner 2001; Wystrach et al. 2011, 2020; Mangan and Webb 2012).

One theoretical mechanism underlying detouring, introduced in Collett et al. (1993, 1998), is that ant (as well as honeybee) navigators can suppress their global vector and retrieve a ‘local vector memory’, a directional and distance estimate of a portion or segment of the full route (Collett et al. 1993, 1998; Srinivasan et al. 1997; Menzel et al. 1998; Collett and Collett 2009, 2015). Local vectors are defined as PI-derived vector segments independent of both the global vector and current vector state. Instead, local vectors are learned and triggered in combination with familiar panoramic views at the site (Collett et al. 1998, 2002; Bisch-Knaden and Wehner 2001; Collett and Collett 2015). Importantly, and perhaps controversially given our current understanding of view alignment, these familiar panorama views trigger the retrieval of local-vector memories rather than directly provide directional information. Yet this presence of the familiar panorama results in many instances in which examples of foragers using a local vector might also be explained through view alignment alone or orientation via motor routine (Baddeley et al. 2011, 2012; Lent 2013; Hoinville and Wehner 2018; Webb 2019; Wolf 2011). This propensity for navigating ants to engage in view matching has made untangling local vectors from view guidance difficult, leading to local vectors being absent in recent PI models (Heinze et al. 2018; Hoinville and Wehner 2018; Stone et al. 2017).

While local vectors as described by Collett and colleagues remain unresolved, recent work in trail-following ants provides evidence of a similar (though mechanistically distinct) phenomenon without the confounding presence of view alignment, with foragers attending only a segment of their global vector during navigation along non-straight-line routes (Plowes et al. 2019; Freas et al. 2020). The most compelling evidence of PI-based navigation to an intermediate site comes from the desert harvester ant, *Veromessor pergandei*. Inbound *V. pergandei* foragers attend only a portion of their global vector while not in contact with the trail pheromone to first reach the pheromone trail rather than directly returning to the nest (Freas et al. 2020).

This behaviour is underpinned by *V. pergandei’s* foraging ecology, a column-and-fan foraging structure. *V. pergandei* individuals initially leave the nest navigating socially along a pheromone column (up to 40m) before the pheromone ends (column head) and foragers fan out alone, off the pheromone, to collect food (Plowes et al. 2013, 2014). Throughout this journey, both in the column and fan, foragers rely on an accumulating PI-derived vector to navigate and do not engage in view alignment (Freas et al. 2019ab; Freas and Spetch 2021). After collecting food, individuals navigate via only a portion of their global vector to first reach the pheromone column before reorienting toward the nest (Freas et al. 2020). The segmentation of the global vector into ‘on’ and ‘off’ pheromone sections is mediated by the pheromone cue, with exposure leading to the re-emergence of the vector accumulated within the column and an orientation change to the global vector direction. In this way the presence of the pheromone triggers a goal shift from the column to the nest after reaching the intermediate goal.

These findings point to the pheromone column functioning as a critical waypoint along the nest-ward route with an olfactory trigger. Yet the underlying mechanism within the PI system is currently unknown as it is unclear if this behavior represents the capacity of multiple vectors running concurrently or if, fitting with local vector theory, portions of the global vector can be neurally suppressed to produce waypoint homing. Interestingly, *V. pergandei* (also see *Formica obscuripes, Freas and Spetch In Press*) also shows evidence of attending to vector segments within non-straight columns, feats which must have a separate triggering mechanism from pheromone presence observed in the fan. Within the pheromone column, when inbound foragers from non-straight-line foraging columns are tested at unfamiliar locations, forager paths do not align with the global vector but only to the current inbound route segment (Freas et al. 2020, Figure 4e; Freas and Spetch *In Press*, Figure 2c). However, untangling orientation to the global vector vs. a vector segment can be difficult due to small directional differences in naturally forming columns.

Here we explore vector-segment way pointing while on the pheromone trail by experimentally creating two-legged column routes to and from food sites, increasing the directional differences between the overall global vector and its underlying route segments and then testing foragers from these routes at a distant, unfamiliar site. We found that foragers navigating along non-straight routes did not home via their global vector and instead navigated via distinct segments of their PI (vector segments), allowing them to directionally retrace separate legs of their inbound route. As this behaviour occurred at a distant site, vector-segment homing was not triggered by familiar external cues, but more likely the navigator’s PI state.

## Methods

### Study Species

For the current study we tested individuals from three *Veromessor pergandei* nests all located within the Verrado Temporary Trails in Buckeye, Arizona, USA (33°19′43.10′′ N, 112°01′02.60′′ W) with nests spread over a 152m area (Nest 1 to Nest 2). Testing with Nest 1 was completed during October to November 2019 while testing with Nest 2 and Nest 3 was conducted during October to November 2021 and March to April of 2022. Across conditions only foragers holding a visible piece of food, cookie pieces provided at the designated feeding site/column head, were tested to ensure homeward motivation. After testing foragers were marked as tested and returned to the nest where they continued to forage over the ensuing days.

### Two-Leg Column

After observing the natural column direction on the previous day, we erected a barrier to force foragers to initially leave the nest in a direction with a 45° discrepancy from their desired foraging direction. Before morning activity commenced at the Nest 1 entrance, low 10cm high barriers were placed on the ground around the nest, extending out for 2m and creating a 30cm wide channel for the foraging column to form traveling away from the nest entrance (Figure 1a; Supplemental Figure 1). This initial segment of the route (nest entrance to the 90° elbow) was designated as Leg 1 of the route. At 2m this channel was left open to allow the foraging column to shift back to the nest’s desired foraging location, leading to a 90° counterclockwise elbow from Leg 1 to Leg 2. Leg 2 extended 2m (with a ∼50cm width) along this new route direction where we placed a food site by spreading crushed cookie pieces on the ground in a ∼15cm diameter area, creating a two-legged column with each leg extending 2m and a 90° elbow between the two legs (Figure 1a). The foraging column ceased extending once reaching this large amount of food with no evidence of foragers moving beyond this point. After collecting food, inbound foragers were observed to retrace this two-legged route, rather than leaving the pheromone column along their global-vector direction to return to the nest directly.

**Figure 1.**
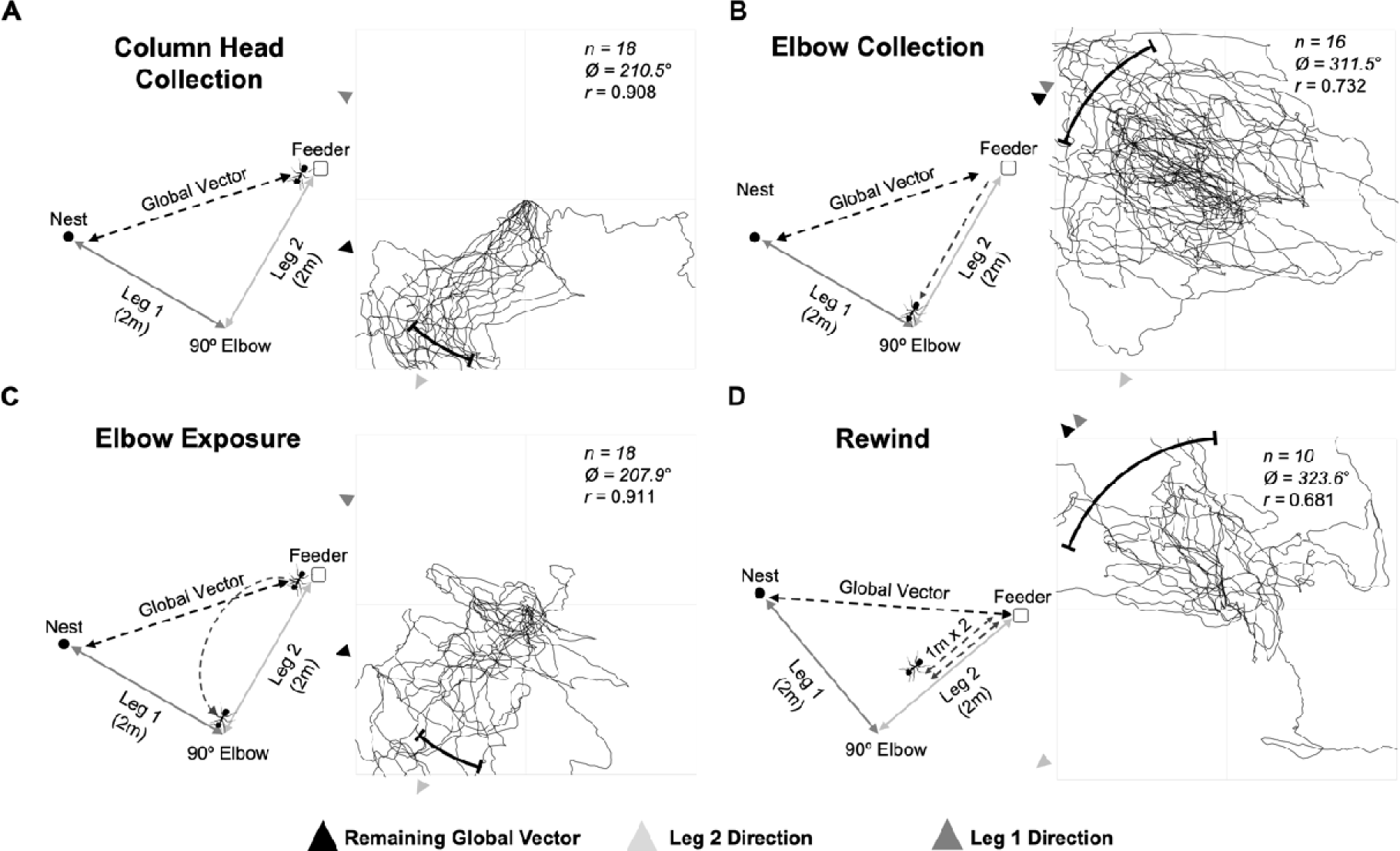
Diagrams and forager paths in the *Two-Leg Column* conditions. (A) Column Head, (B) Elbow Collection, (C) Elbow Exposure, and (D) Rewind conditions. Forager paths extend until grid exit and statistics indicate 1m heading directions. Black arrows: foragers’ remaining PI derived global vector; Grey arrows: Leg 2 compass direction; Dark grey arrows: Leg 1 compass direction; Black bars: 95% CI of headings. n, number of individuals; Ø, mean vector; r, mean vector length.

Foragers were initially randomly assigned one of three separate conditions, collected and displaced to a distant unfamiliar location (>100m from all nests) that was flat and devoid of vegetation. At this site, a 2m × 2m grid of 1m squares was erected using metal pegs and string. Foragers were released at the centre of this grid and their paths were recorded until they left the grid using a HD GoPro camera (30fps and 3840 × 2160 pixels image size) which was positioned 1.5m above the release point facing straight down. In the first condition, foragers (n=18) were collected just as they grabbed a cookie piece from the column head/food site (*Column Head* condition; Figure 1a). In the second condition, foragers (n=16) were allowed to travel, with food, back along Leg 2 to the 90° elbow (*Elbow Collection* condition; Figure 1b) and were collected in the last 10cm of Leg 2 just as they reached the barrier that defined the start of Leg 1. In the third condition, foragers (n=18) were collected from the column head and released at the 90° elbow for 20sec (*Elbow Exposure* condition; Figure 1c). After this exposure period, foragers were re-collected and displaced to the distant site.

These three conditions were completed over two consecutive days while the column structure and direction remained stable. On the third day, the morning column formation shifted 20° counterclockwise. We re-erected the barriers to create the same two-legged column with a 90° counter-clockwise elbow and conducted a final condition (*Rewind* conditio*n;* Figure 1d) where foragers (n=10) were ‘rewound’ along Leg 2. Rewinding along a route allows us to update the navigator’s path integrator state (to the 90° elbow) without exposing the individual to the external cues at this site, excluding them as triggers of an orientation shift. Foragers with food were allowed to travel halfway back (1m) along the 2^nd^ Leg where they were collected and released back at the column head to repeat the first 1m of Leg 2 again. After 1m these individuals were again collected and tested distantly.

### Two-Leg Column - Full paths

#### Nest 1

As we observed evidence of foragers in the *Column Head* and *Elbow Exposure* conditions turning away from their global vector and orienting instead only along the Leg 2 compass direction towards the 90° Elbow, we chose to further explore this behaviour by conducting a replicate of the *Column Head* condition with a separate set of foragers (n=12) while recording the full forager paths beyond 1m. Around Nest 1, the barrier along Leg 1 was extended 2.5m with a 30cm width, while Leg 2 remained at 2m with the column width measuring 50cm (Figure 2a; Supplemental Figure 1). Additionally, we expanded the grid at the testing site to a 5m × 5m grid of 1m squares and again collected foragers from the column head/food site, just as they grabbed a food piece, and released them at the distant site. Forager paths were again recorded with an overhead camera for 5min or until they left the grid.

**Figure 2.**
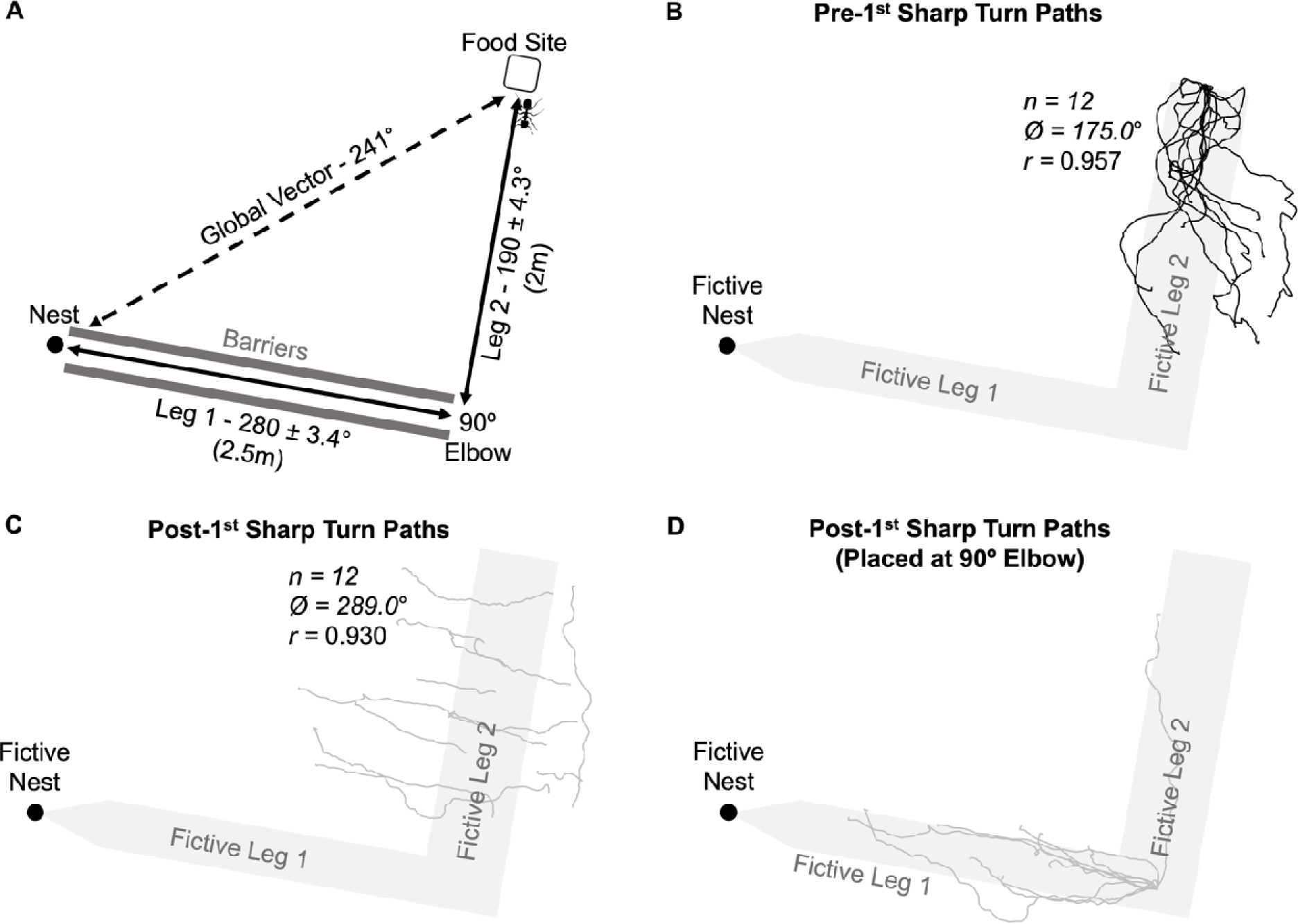
Diagram and forager paths in the Nest 1 *Full Path* condition (distant site). The fictive column (Leg 1 and Leg 2) is provided in grey to illustrate forager paths alignment with the inbound route. (**A**) Foraging column with Legs 1 and 2 separated by a 90° Elbow. All foragers were collected from the column head/ food site and displaced distantly. (**B**) Shows 12 forager paths (black) until their first sharp (>60°) turn at the distant site. (**C**) Shows the post first sharp turn paths (grey) of these same 12 foragers until they conduct a second sharp turn. After this segment foragers began to search. (**D**) As foragers in all conditions underestimated the distance of Leg 2, we illustrated foragers’ post sharp turn directional alignment with the Leg 1 compass direction by placing the post turn origin as if it had occurred at the 1^st^ Elbow. n, number of individuals; Ø, mean vector; r, mean vector length.

When collecting path data, typically the end of navigation via path integration and the onset of search is defined as the first sharpL(> 50-60°) turn with the forager continuing in the new direction for at least 20cm (Schwarz et al. 2017; Freas et al. 2020, 2021). As these new, post-turn orientations are typically non-straight and uniform in direction, denoting the onset of the systematic search spiral, they are often excluded from analysis. For this experiment, we chose to examine path headings before and after each forager’s first sharp turn > 60° (consistent with Freas et al. 2020) to characterize if foragers began to search after this turn or if orientation was altered to a new predicted direction (global vector or Leg 1 compass direction).

#### Nest 2

After testing with Nest 1 foragers, we conducted a full path replicate using a second nest (Nest 2). Here the natural nest heading curved clockwise so that the column passed between two bushes, located 3m from the nest. We established a 90° elbow two-legged foraging column (Figure 3a; Supplemental Figure 1) using the same procedure as previous conditions with a 3m long, 50cm wide Leg 1 with a 90° clockwise elbow and a 2.5m long, 40cm wide Leg 2 (column width of the second leg extending from Nest 2 was constrained by vegetation). Foragers (n=14) were collected at the column head, just as they grabbed food, and displaced to the distant site where their inbound paths were recorded identically to previous conditions.

**Figure 3.**
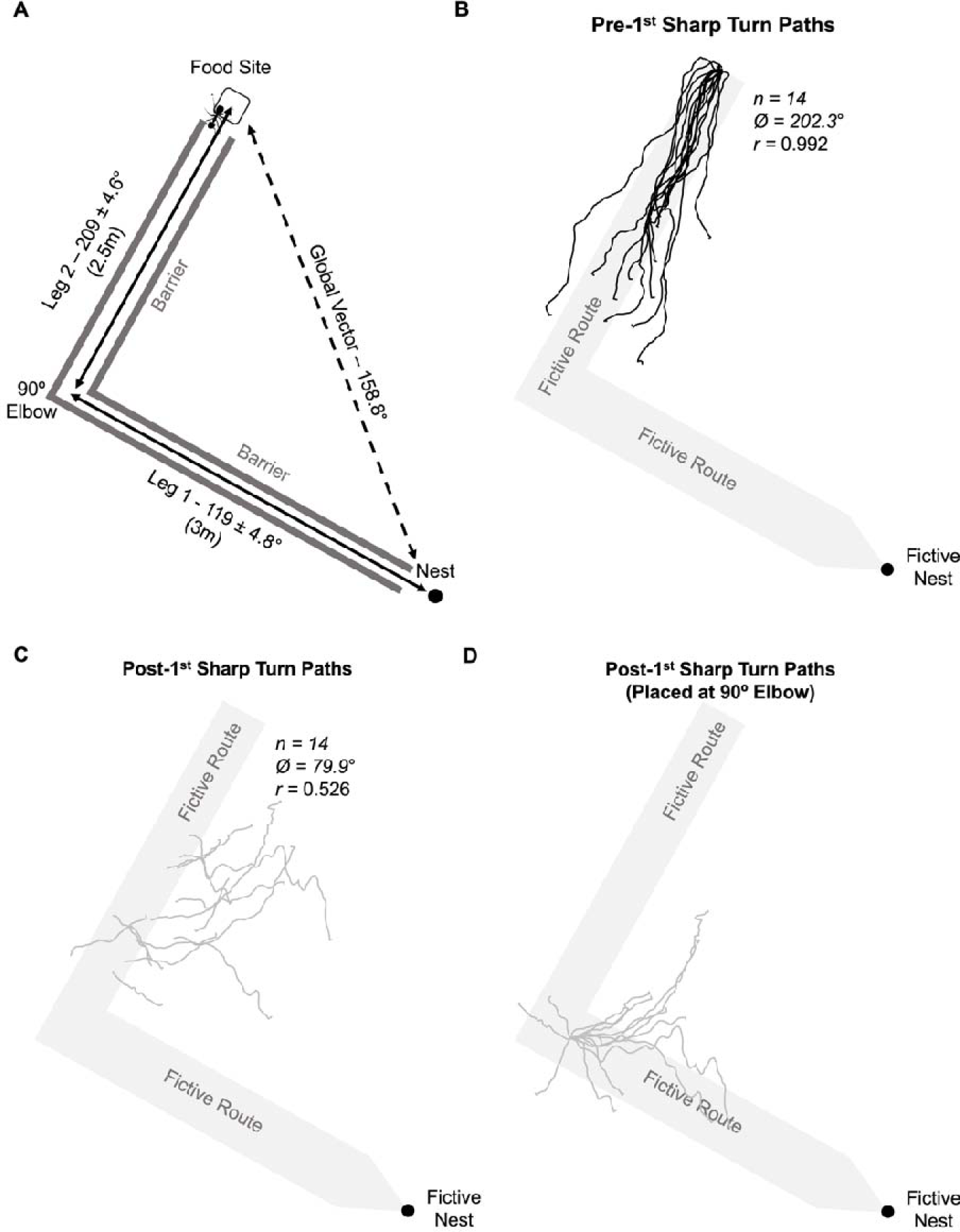
Diagram and forager paths in the Nest 2 *Two-Leg Column Full Path* condition (distant site). The fictive column (Leg 1 and Leg 2) is provided in grey to illustrate forager paths alignment with the inbound route. (**A**) Foraging column created with Legs 1 and 2 separated by a 90° turn. (**B**) 12 forager paths (black) until their first (>60°) turn. (**C**) Post-turn paths (grey) until foragers conduct a second sharp turn (search onset). (**D**) Given observed underestimation of Leg 2, we illustrated foragers’ post sharp turn directional alignment with the Leg 1 compass direction by placing the post turn origin as if it had occurred at the 1^st^ Elbow. n, number of individuals; Ø, mean vector; r, mean vector length.

### Three-Legged Column - Full paths

#### Nest 2

We next characterized how non-nest orientation (along Leg 2) was initiated by collecting foragers moving along their global-vector direction, before their pheromone column shifted away from this direction. Around Nest 2 we erected barriers creating a three-legged column (Figure 4a). Leg 1 of the column extended 3m from the nest entrance (width 50cm) and ended with a 90° counterclockwise elbow (1^st^ Elbow) and a 3m Leg 2 (width 40cm). At the end of Leg 2 the arena shifted 45° clockwise (2^nd^ Elbow) with a final Leg 3 extending 0.5m where we designated a food site and the column ended. This area setup resulted in inbound foragers traveling along Leg 3 traveling in their global vector before the pheromone column turns away from this direction along Leg 2. As in previous tests, foragers (n=11) were collected just as they grabbed a food piece and transferred to the distant site, where their paths were recorded using an overhead camera for 5min (or grid exit). As foragers’ inbound paths from the food site along the column initially allowed foragers to orient along a leg (Leg 3) which was directionally identical to the forager’s global vector for 50cm, at the distant site we collected initial headings at 30cm as well as headings pre and post the first sharp turn, to catalogue any observable initial orientation to this global vector and an orientation change to the Leg 2 segment.

**Figure 4.**
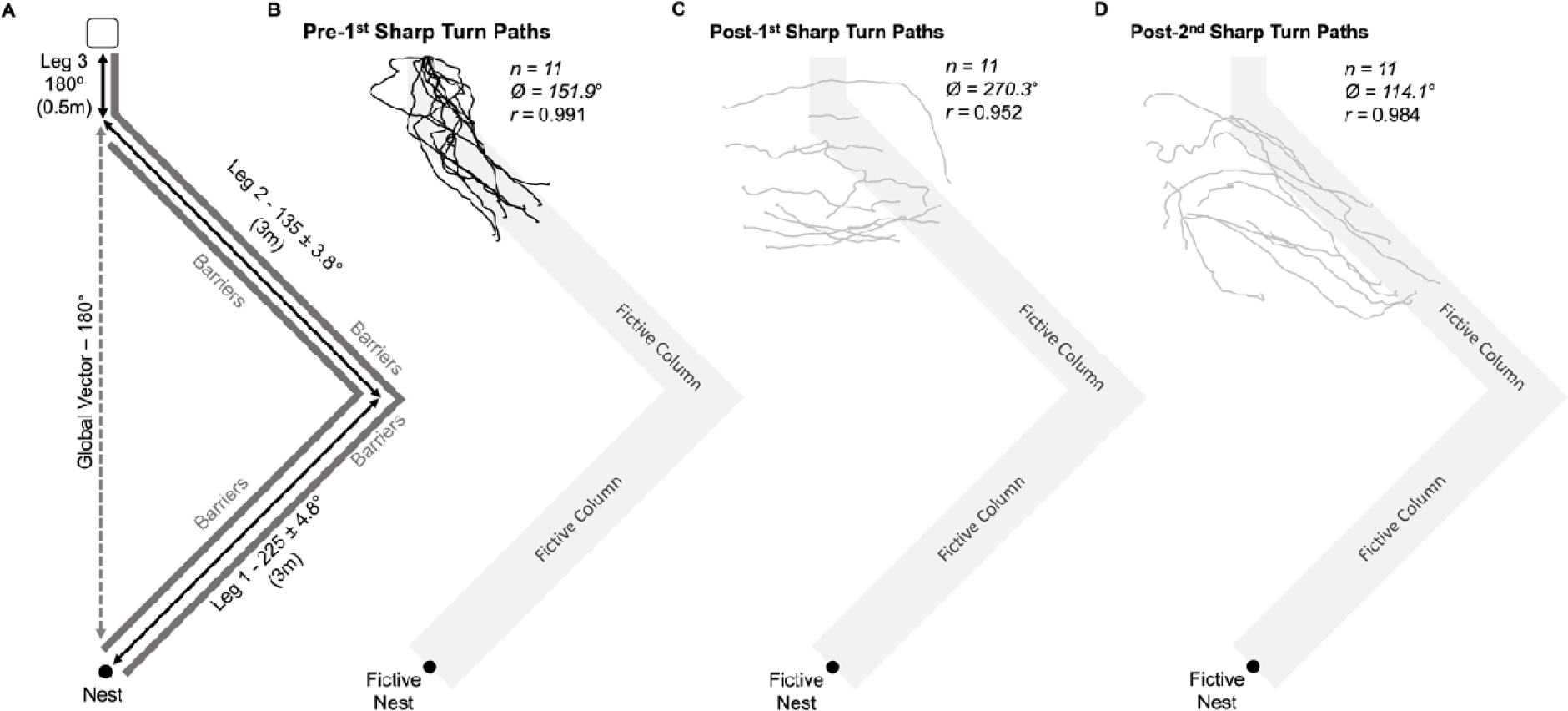
Diagram and forager paths in the *Three-Leg Column* condition (Nest 2). The fictive column (Leg 1, 2 and 3) is provided in grey to illustrate forager paths alignment with the inbound route. (**A**) Foraging column with Legs 1 and 2 separated by a 90° 1^st^ Elbow, while Leg 2 and 3 separated by a 45° 2^nd^ Elbow. (**B**) Forager paths (black) until their first sharp (>60°) turn. (**C**) Post first turn paths (grey) of same foragers until a second sharp turn. (**D**) Shows the third path segment after each forager’s second sharp turn (headings remained directed).

#### Nest 3

Testing at Nest 3 was conducted to further explore the limits of foragers’ ability to suppress vector segments. We observed some evidence with Nest 2 of foragers initially orienting to Leg 3, which could be explained as individuals ignoring two segments of their global vector. However, given Leg 3 was in the global-vector direction, an alternative explanation is that vector segment use only occurs once foragers turn away from this global vector direction and orient along Leg 2. This would result in only a single waypoint along the homeward route and may represent a limiting factor for the path integrator system to follow vector segments within the column. With Nest 3 we altered the three-legged set up to reduce these confounds by having the Leg 3 diverge from the global vector by 36° (Figure 5a; Supplemental Figure 1).

**Figure 5.**
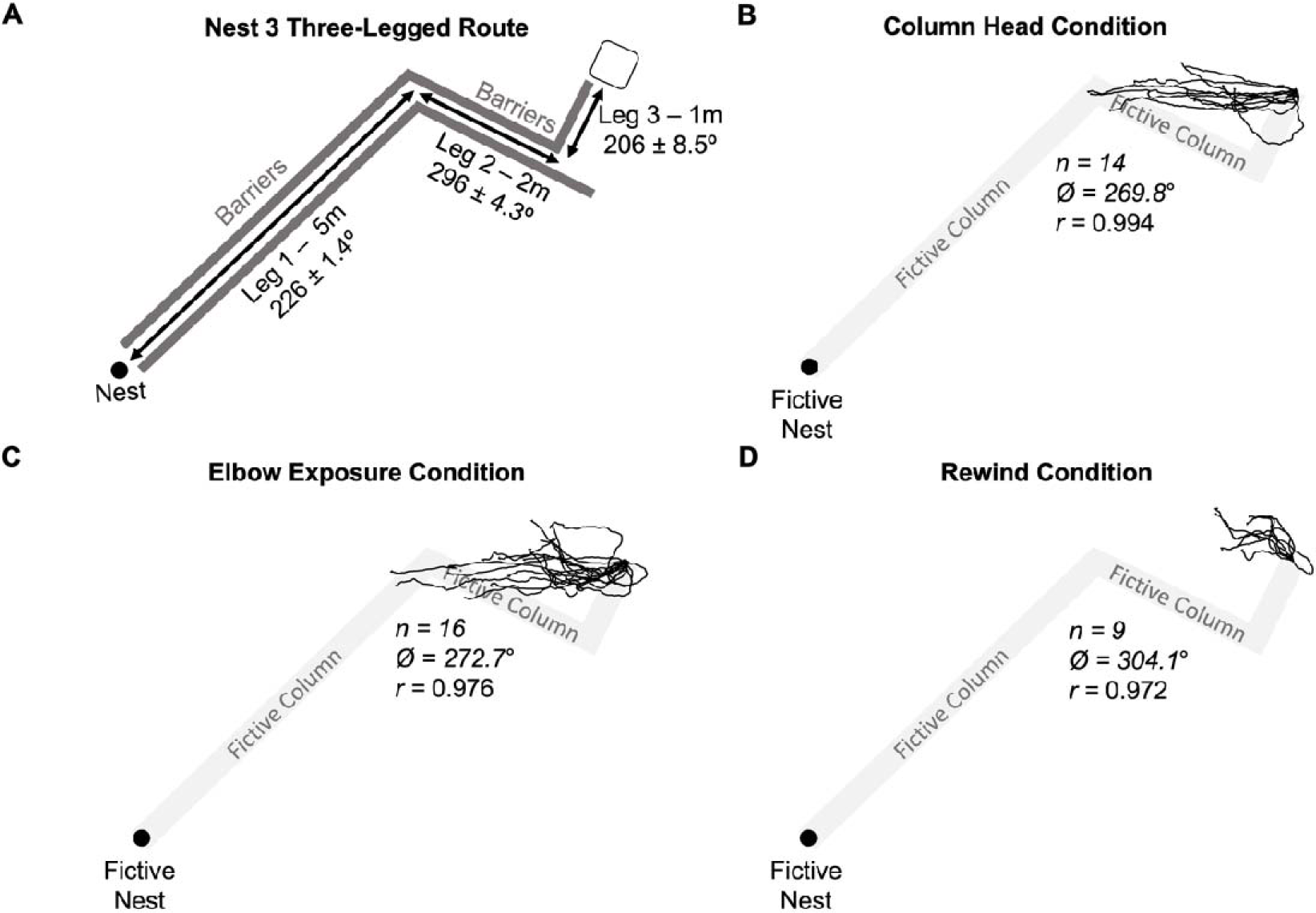
Diagram and forager paths at the distant site in the *Three-Leg Column* conditions for Nest 3. The fictive column segments of Leg 1, Leg 2, and Leg 3 are illustrated in grey to allow for comparisons of how forager paths align with the inbound column directions and distances. (**A**) Nest 3 foraging column where Legs 1 and 2 were separated by a 70° 1^st^ Elbow and Leg 2 and 3 were separated by a 90° 2^nd^ Elbow. (**B**) Column Head forager paths (black) until their first sharp (>60°) turn. (**C**) Turn Exposure forager paths until their first sharp (>60°) turn. (**D**) Rewind condition forager paths until their first sharp (>60°) turn.

We erected an arena with three legs. Leg 1 extended 6m (30cm width) before turning 70° clockwise (1^st^ Elbow) and extending 2m along Leg 2 (30cm width). At the end of Leg 2 the arena turned 90° counterclockwise (2^nd^ Elbow) and extended 1m along Leg 3 (30cm width), where we placed a food site to end the foraging column. This created a column where inbound foragers after collecting food must first orient 32° to the left of their global vector (along Leg 3) then turn 54° to the right of their global vector to follow along Leg 2 of the column. Foragers were collected and randomly assigned one of three separate conditions. In the *Column Head* condition, foragers (n=14) were collected just as they grabbed a food piece from the column head at the end of Leg 3. In the second condition, foragers (n=16) were collected from the column head and released at the 70° 1^st^ Elbow between Leg 1 and Leg 2 for 20sec before being recollected and tested distantly (*Elbow Exposure*). In the final condition, foragers (n=9) were rewound along Leg 3 to set their vector state as if they were at the 2^nd^ Elbow but without being exposed to the cues at this site. Foragers were allowed to travel halfway back (50cm) along Leg 3, collected and released back at the column head and then re-collected at the halfway point and tested distantly (*Rewind Condition*). Forager paths were recorded at the distant site under the same methodology as previous conditions.

### Statistical Analysis

Video analysis was conducted to digitize forager paths using GraphClick. Forager positions from release were recorded by taking their position in 0.2-sec increments. Heading directions in each condition were first analyzed using Oriana software to determine if they were directed using Rayleigh Tests for circular data (Fisher 1993, equations 4.17 & 4.18). If headings were considered directed via the Rayleigh Test (α = 0.05), we further analyzed if the mean direction of observed headings was in a particular predicted direction using the 95% confidence interval around the mean of foragers headings. Confidence intervals were calculated through the standard error of the mean heading direction based on the mean vector length (Fisher 1993, equation 4.42). When deciding the compass direction of individual legs of the column, we chose the range of headings representing the width of the column rather than the specific centre point in order to account for columns forming tightly around inner bends of each elbow (some columns formed hugging the barrier around elbows). In the three-legged route conditions where we also analyzed orientation to the 1^st^ Elbow, we took compass measurements from the food site to the inner edge of this Elbow to compare with heading data. Between-condition comparisons of mean heading directions were conducted using Watson-Williams F-Tests (Fisher, 1993, p.126; Batschelet, 1981, p.95) while paired comparisons within the paths of individual foragers (i.e., headings pre/post-turn) were analyzed using Moore’s Paired Tests (Zar 1999). When multiple comparisons were made with the same condition, Bonferroni corrections (^α^*/_n_*) were used to determine significance level (α = 0.025).

## Results

### Two-Leg Column

*Column Head* forager paths were significantly directed at 1m (Rayleigh Test; *Z* = 14.83; *p* < 0.001; Figure 1a) and only the compass direction of Leg 2 of the column (210±7.1°) was within the 95% CI of headings (210.5±5.9°), while both the compass directions of the global vector (255°) and the Leg 1 column (300±4.3°) were well outside this range, signifying orientation not to the nest but along the (Leg 2) route to the fictive position of the 90° Elbow. This elbow orientation remained consistent for forager headings in the *Elbow Exposure* condition (207.9±5.8°) which were also significantly directed (Rayleigh Test; *Z* = 14.94; *p* < 0.001; Figure 1b) only in the compass direction of Leg 2 (210±7.1°) towards the 90° Elbow (95% CI) while both the global vector (255°) and the Leg 1 (300±4.3°) compass directions were well outside this range. Mean heading directions between these conditions (*Column Head vs. Elbow Exposure*) did not significantly differ (Watson-Williams F-Test; *F*_(1,34)_ = 0.09; *p* = 0.764) suggesting exposure to the external cues present at the Elbow did not influence foragers’ compass headings.

In contrast, the headings of foragers in the *Elbow Collection* condition (311.4±11.2°) were significantly directed (Rayleigh Test; *Z* = 8.56; *p* < 0.001; Figure 1c), yet headings were not directed in the compass direction of Leg 2 (210±7.1°). Instead, Leg 1’s compass direction at 300(±4.3°), which was also the direction of foragers’ remaining global vector, was within the 95% CI of headings. Foragers in the *Rewind* condition (323.6±16.4°) were similarly directed (Rayleigh Test; *Z* = 4.64; *p* = 0.006; Figure 1d) in the Leg 1 compass direction (320±4.3°) and remaining global vector (320°) while not in the Leg 2 compass direction at 230±7.1°. Mean heading directions between these conditions did not significantly differ between the *Rewind* and *Elbow Collection* conditions (Watson-Williams F-Test; *F*_(1,24)_=0.36; *p*=0.552), indicating that the change in headings was not triggered by exposure to external cues present at the Elbow.

### Two-Leg Column - Full paths

#### Nest 1

Foragers exhibited pre-turn headings that were directed (Rayleigh Test; *Z* = 10.98; *p* < 0.001; Figure 2b) with only the compass direction of the Leg 2 column (190±7.1°) within the 95% CI of headings (175.0±11.0°). Both the global vector (241.3°) and Leg 1 (280±3.4°) were well outside this heading range. The average distance travelled pre-turn (µ±s.e.m. = 1.14±0.13m) was well short of the 2m distance to the 1^st^ Elbow (57%).

Post-turn headings significantly shifted away from the pre-turn direction (Moore’s Paired Test; *R*_(12)_ = 1.80; *p* < 0.001). Post-turn headings (289.0±14.1°) remained directed (Rayleigh Test; *Z* = 10.38; *p* < 0.001; Figure 2c) with only Leg 1‘s compass direction (280±3.4°; Figure 2c,d) within the 95% CI of headings. When we individually calculated each forager’s remaining global vector compass direction at the turn position, this direction (260.7±5.0°) was significantly different from the observed post-turn headings (Moore’s Paired Test; R_(12)_=1.802; p<0.001) with headings biased on average 30.57±13.3° to the right of the remaining global vector. Mean distance travelled during this second path segment (µ±s.e.m.=0.90±0.13m) was short of Leg 1’s 2.5m distance (36%) before the ants conducted another sharp turn. Beyond this second turn, forager paths were uniform and no longer directed (Rayleigh Test; *p* > 0.05), suggesting the onset of search.

#### Nest 2

Foragers headings were directed (Rayleigh Test; *Z* = 10.98; *p* < 0.001; Figure 3b) and once again only Leg 2 (209±4.6°) was within the 95% CI of headings (202.3±4.4°), while both the compass directions of the global vector (158.8°) and Leg 1 (119±4.8°) were well outside this range. Again, foragers did not travel the full 2.5m Leg 2 distance before turning (µ±s.e.m. = 1.87±0.12m), with foragers completing 74.7% of the true distance. After the first turn, foragers’ headings significantly shifted away from the pre-turn headings (Moore’s Paired Test; *R*_(14)_ = 1.81; *p* < 0.001) and these new headings remained directed (Rayleigh Test; *Z* = 3.87; *p* = 0.018; Figure 3c,d), suggesting foragers had not begun to search. Only the Leg 1 compass direction (119±4.8°) was within the 95% CI of these headings (79.9±39.2°) while Leg 2 (209±4.6°) was well outside this range. We again calculated each forager’s remaining global vector compass direction from their position at their first sharp turn and this compass direction (139.9±4.4°) was significantly different from the observed headings post-turn (Moore’s Paired Test; *R*_(14)_ = 1.64; *p* < 0.001) with observed mean headings 59.83±35.8° to the left of the global vector direction. Post-turn foragers were oriented only to Leg 1 of their vector and not to their remaining global vector. After the second > 60° turn, mean path headings were uniform and no longer directed (Rayleigh Test; *Z* = 1.37; *p* = 0.257), indicative of the onset of search.

### Three-Legged Column - Full paths

#### Nest 2

Initial headings (30cm) were directed (Rayleigh Test; *Z* = 7.48; *p* < 0.001; Figure 4a) and only the compass direction of the global vector (180°) was within the 95% CI of these headings (165.8±24.2°) while the compass direction of the Leg 2 column (135±3.8°) was just outside this range (Leg 1 at 225±4.8° was well outside). However, the compass direction from release to the 1^st^ Elbow at 149°, representing a combination of the Leg 3 and Leg 2 columns, was within the 95% CI of initial headings at 30cm. After foragers’ first sharp turn, headings remained directed (Rayleigh Test; *Z* = 10.81; *p* < 0.001; Figure 4a) but had shifted from their initial orientation counterclockwise by 13.9° towards Leg 2 (though this change was not significant; Moore’s Paired Test; *R*_(11)_ = 0.987; *p* > 0.05). Interestingly, the global-vector compass direction (180°) was no longer within the 95% CI of headings of foragers at their first turn (151.9±5.2°). Leg 2’s compass direction (135±3.8°) also fell outside this range, yet the 1^st^ Elbow direction (149°) was within the 95% CI suggesting orientation to a combination of Leg 2 and 3 (to 1^st^ Elbow), representing a potential limitation to the system.

Just as in previous conditions, foragers again underestimated the distance to the 1^st^ Elbow, conducting their first turn well before the true distance (µ±s.e.m. = 1.34±0.10m) at 40.6% of the distance (3.3m). Post first turn, forager headings significantly shifted away from the pre-turn headings (Moore’s Paired Test; *R*_(11)_ =1.80 ; *p* < 0.001) and headings remained directed (Rayleigh Test; *Z* = 9.98; *p* < 0.001; Figure 4c). Despite highly directed headings (µ±95% CI = 270.3±12.3°), no predicted compass direction fell within the 95% CI (Global Vector: 180°; Leg 2: 135±3.8°; Leg 1: 225±4.8°) and these paths extended for 1.11±0.17m.

Unlike previous conditions, after foragers made their second sharp turn, post-turn headings were still directed (Rayleigh Test; *Z* = 10.65; *p* < 0.001; Figure 4d), suggesting foragers were not initiating search. Yet, headings during this third path segment were (µ±95% CI = 114.1±7.1°) well to the left of the global-vector direction (180°) and neither the Leg 2 (135±4.6°) nor Leg 1 (225±4.8°) column directions were within this range. When we calculated the compass direction of the 1^st^ Elbow from the position of each forager’s second turn (123.7±8.2°) and compared this to each forager’s heading (116.8±9.3°), they were not significantly different (Moore’s Paired Test; *R*_(11)_ = 0.769; *p* > 0.05). In contrast, these headings were significantly different from each forager’s remaining global vector at the onset of the 3^rd^ path segment (Moore’s Paired Test; *R*_(11)_ = 1.80; *p* < 0.001) with headings on average 59±7.1° to the left of the remaining global vector suggesting foragers were homing to the 1^st^ Elbow, not the nest. By the end of this path segment, the 5min recording period expired for four of the eleven ants, making further analysis of the remaining seven ants after this point difficult.

#### Nest 3

In all conditions, foragers exhibited directed initial headings (*Column Head* µ±95% CI = 245.9±15.4°, Rayleigh Test; *Z* = 11.36; *p* < 0.001; *Elbow Exposure* µ±95% CI = 224.3±19.3°, Rayleigh Test; *Z* = 9.85; *p* < 0.001; *Rewind* µ±95% CI = 318.9±15.9°, Rayleigh Test; *Z* = 7.94; *p* < 0.001; Figure 5). Initial mean headings in *Elbow Exposure* did not significantly differ from *Column Head* foragers (Watson-Williams F-Test; *F*_(1,28)_ = 2.87; *p* = 0.101). In contrast, initial mean headings in the *Rewind* condition were significantly different from *Column Head* foragers (Watson-Williams F-Test; *F*_(1,21)_ = 45.36; *p* < 0.001). In *Column Head* foragers, only the global vector (243°) fell within the 95% CI of initial headings while in the *Elbow Exposure* condition, both the global-vector compass direction (243°) and Leg 3 column compass direction (206±8.5°) were within the 95% CI of headings. In contrast, no predicted compass direction was within the 95% CI of *Rewind* headings, though the compass direction of the Leg 2 column direction was only 2.7° outside this range at 296±4.3°.

By their first turn, foragers in all conditions remained oriented (*Column Head:* µ±95% CI = 269.8±3.8°, Rayleigh Test; *Z* = 13.83; *p* < 0.001; *Elbow Exposure:* µ±95% CI = 272.7±6.2°, Rayleigh Test; *Z* = 15.24; *p* < 0.001; *Rewind:* µ±95% CI = 304.1±10.6°, Rayleigh Test; *Z* = 8.51; *p* < 0.001). Just as when comparing initial headings between conditions, there was no significant difference in mean heading direction between the *Column Head* and *Elbow Exposure* conditions (Watson-Williams F-Test; *F*_(1,28)_ = 0.557; *p* = 0.462), suggesting exposure to the cues at the 1^st^ Elbow did not influence forager headings, and *Rewind* foragers, consistent with a PI state placing them at the 2^nd^ Elbow, exhibited mean headings that remained significantly different from *Column Head* foragers (Watson-Williams F-Test; *F*_(1,21)_ = 60.52; *p* < 0.001).

In both *Column Head* and *Elbow Exposure* conditions, headings by foragers’ first turn had shifted significantly from initial heading directions, 23.9° clockwise (Moore’s Paired Test; *R*_(14)_ = 1.865; *p* < 0.001) and 48.4° clockwise, respectively (Moore’s Paired Test; *R*_(16)_ = 1.847; *p* < 0.001), while *Rewind* foragers exhibited no significant directional shift (Moore’s Paired Test; *R*_(9)_ = 0.763; *p* > 0.05). Headings at the first sharp turn in *Column Head* and *Elbow Exposure* conditions were no longer directed towards the global-vector direction (243° outside 95% CI) and as we observed during testing with the Nest 2 three-legged route, no individual leg compass direction was within the observed 95% CI of headings. However, the compass direction from release to the 1^st^ Elbow at 268°, representing a combination of Legs 2 and 3, was within the 95% CI of headings in both conditions. In *Rewind* foragers, only the compass direction of Leg 2 (296±4.3°) was within the 95% CI of headings, consistent with the rewinding manipulation creating a PI state at the 2^nd^ Elbow. *Column Head* foragers travelled 80.8% (µ±s.e.m. = 1.81±0.17m) and *Elbow Exposure* foragers travelled 71.8% (µ±s.e.m. = 1.61±0.16m) of the (2.24m) actual distance to the 1^st^ Elbow before turning. *Rewind* foragers only travelled 41.3% (µ±s.e.m. = 0.83±0.07m) of the actual 2m Leg 2 distance before turning.

After foragers’ first turn in all three conditions, paths remained directed (*Column Head,* µ±95% CI = 111.0±45.8°, Rayleigh Test; *Z* = 3.06; *p* = 0.044; *Elbow Exposure,* µ±95% CI = 105.4±18.7°, Rayleigh Test; *Z* = 10.17; *p* < 0.001; *Rewind,* µ±95% CI = 118.3±26.1°, Rayleigh Test; *Z* = 6.41; *p* = 0.003); in all of these conditions, however, we observed no evidence of orientation to Leg 1 or any other predicted inbound direction. Foragers were instead oriented back in the direction of the release point (88° in the *Column Head* and *Elbow Exposure*; 116° in the *Rewind* condition), suggesting foragers had abandoned inbound orientation and are searching/backtracking to re-enter the pheromone.

## Discussion

Within the field of ant navigation, the ability for individuals to home to intermediate sites along the inbound route instead of their origin is controversial. Two strategies which are comprehensively modelled, view-based memory alignment and path integration (PI), inbound navigators are thought to home specifically to the nest and lack evidence of inbound homing to specific non-nest locations even when detouring. PI is currently conceptualized and modelled as a singular updating global estimate of the navigator’s start location (nest) in relation to their current position. Theories, such as local vectors, which propose the ability to rely upon independent portions of the global PI to produce detour behaviours, have largely been abandoned, with these behaviours now believed to be performed via view alignment, motor routines and reinforcement learning (Wolf 2011; Hoinville and Wehner 2018; Stone et al. 2017; Webb et al. 2019; Wystrach et al. 2020). Yet, our work on fan and column foraging ants, shows that inbound ant navigators possess the ability to attend to a portion of their global vector to first reach a waypoint along the inbound route before reorienting to the nest. These PI-based waypoints indicate a needed expansion of the path integration system beyond estimating the start-point to include PI’s ability to waypoint along the inbound route. Trail-following *V. pergandei* are known to accumulate PI estimates both while in their pheromone-marked column and after they leave this column to search for food alone in a foraging fan. Fan foragers first navigate back to the pheromone column before re-orienting to the nest, attending to only the portion of their PI accumulated while off the column while the rest of the global vector appears absent from their orientation. Successfully reaching this column waypoint is critical for the column portion of the vector to be re-expressed, facilitating the return along the pheromone column to the nest. This behavioural evidence indicates an olfaction-mediated distinction between PI accumulation in the presence and absence of the pheromone, allowing foragers to either: 1.) retain a second vector which begins at the column head as they leave the pheromone or 2.) the ability to home via part of their global PI estimate to reach intermediate sites along the route. Here we show this ability also exists within the pheromone column itself, with foragers showing evidence they attend to only a segment of their accumulated vector within the column, presumably for the purpose of retracing curved routes while maintaining contact with the pheromone during inbound navigation.

Across all conditions, foragers oriented along only part of their global-vector estimate towards a waypoint along the inbound pheromone trail (the elbow). Foragers in two-legged routes with distinct segments separated directionally by 90° oriented to only the second segment (Leg 2), heading towards the elbow instead of the nest. When inbound foragers were exposed to the cues present at this elbow before testing, while their PI state remained unchanged, forager paths did not update to their global vector or Leg 1, instead continuing to orient via only the Leg 2 segment. When foragers’ PI state was updated to the elbow, either through elbow collection or rewinding, headings changed to become directed along Leg 1 to the nest. Together, these headings indicate that foragers travel along portions or vector segments of their global PI and that the cues (both visual and olfactory) present at the elbow does not trigger this reorientation. Instead, the forager’s current PI state underlies the orientation change from attending only Leg 2 to reorienting to Leg 1 of the route. Only after running off most of the Leg 2 portion of the route did foragers re-orient to Leg 1.

Our pre/post sharp turn path analysis provides further evidence that the orientation switch from Leg 2 occurs in the absence of any familiar cues. Foragers tested distantly oriented along Leg 2 and not their global vector before conducting a sharp turn (albeit underestimating the correct distance to the elbow), after which foragers oriented along the Leg 1 of the inbound route and not their remaining global vector. As this directional change occurred at an unfamiliar location, it is doubtful terrestrial-view-based cues triggered the observed orientation change. Instead, after the forager runs off most its Leg 2 vector segment, it reorients to Leg 1.

### Local vectors?

The observed vector segment orientation both in the fan and within the column during route retracing, is evidence of PI-based homing to a waypoint along the inbound route. Such a system would share similarities to the local-vector theory developed in solitarily foraging, non-pheromone-using ants (Collett et al. 1998). In these solitarily foraging species, local-vector theory posits that familiar panoramas trigger the retrieval of a PI-based memory of a vector segment, independent of directional information from the panorama, global vector and current vector state (Collett and Collett 2015). Here, *V. pergandei* foragers rely on PI-based vector segments in order to home to locations along the route, first to re-enter the pheromone column and then to retrace the inbound column. Attending to these segments is independent from panorama cues, being instead triggered by a non-directional olfactory cue (the pheromone) or by the PI-state, thereby removing the view-alignment confound for local vectors in solitarily foraging species (Baddeley et al. 2011, 2012; Collett and Collett 2015; Webb 2019). Vector segments in *V. pergandei* do not occur independent of the navigator’s global vector and current PI state. In fan foragers, the column portion of the global vector is not represented in forager headings while off the pheromone but once a forager contacts the pheromone again, any remaining fan vector is combined with the column vector (Freas et al. 2020). On the column, foragers attended the Leg 2 vector portion of their foraging route before reorienting to the Leg 1 direction after finishing most of their Leg 2 route. In both cases, *V. pergandei’s* current PI state plays a clear role in orientation, making vector segment homing distinct from local vector theory.

### Multiple vectors or vector suppression?

PI-based waypoints are in clear contradiction with PI as a singular memory estimate of the nest and points the PI system’s ability to waypoint to intermediate goals along the inbound route. Importantly, current models of path integration in insects do not account for way-pointing (Heinze et al. 2018; Hoinville and Wehner 2018; Stone et al. 2017; Webb et al. 2019), suggesting an expansion of the capabilities of the path integration system is warranted. Such an expansion would require untangling how segmenting portions of the outbound route could be articulated. We propose two possibilities. First, navigators may be able to retain multiple accumulating vectors at once during their foraging trips which can be weighted differentially along the inbound foraging route to produce way-pointing. There is some evidence that multiple vector memories is possible in Hymenoptera with honeybees possessing two distinct odometer memories, one to communicate to nest mates, and one personal (Dacke and Srinivasan 2008). In ants, the potential for two vector memories based on separate distance estimates from optic flow and the pedometer has also been posited (Wolf et al. 2018). A similar phenomenon may occur in *V. pergandei* where one vector estimate begins at the nest and a second vector estimate accumulation begins running in the absence of the pheromone cues as the forager leaves the column (fan vector). A second theory for vector segment behaivours would take inspiration from local-vector theory via the suppression of the global vector in favor of a vector segment of the route (Collett et al. 1993, 1998). Here, the ant would accumulate a single vector estimate, yet this global vector would be partially suppressed in order to way-point to intermediate goals along the inbound route. Yet in both models, how separate vector segments within the column would be triggered remains unclear (possibly the large rotations in the overhead celestial compass or the motor routine which occurs at the elbow sites). Under both models, evidence from Freas et al. (2020; Figure 4e) and the shortcutting of ants in three legged routes for Nest 2 and Nest 3 suggests *V. pergandei* foragers might be limited to a single waypoint within the column.

A second mechanistic question concerns the triggering mechanism within the column. While separating and reintegrating fan and column vector memories clearly involves an olfactory input, vector memories within the column cannot be triggered by the pheromone presence. Additionally, other aspects of the pheromone column, including deposit intensity are unlikely to be the trigger as elbow exposure did not elicit a heading change. Instead, it appears that the forager must first run down a portion of its current vector segment before choosing to reorient along another leg. It is likely the pheromone still plays some roll in this re-orientation decision as observed by the early turns when testing occurred in the pheromone’s absence. In contrast, foragers observed reorienting on the column, were highly accurate at re-orienting at the 1^st^ Elbow. This suggests that foragers tested while in contact with the pheromone (acting as a reassurance cue: Wetterer et al. 1992; Czaczkes et al. 2011; Freas and Spetch 2021) would continue to run off its vector segment to completion before re-orienting. While running off a vector segment, the forager is likely also continuously attending the pheromone’s presence along its chosen route. Prolonged periods of pheromone absence likely increase uncertainty and early abandonment of the current heading, just as we see in foragers from straight columns abandoning their global vector early (Freas et al. 2020; Freas and Spetch 2021). Therefore, despite the vector segment switch being triggered by the forager’s PI, the pheromone’s presence/absence still influences how far foragers attend to each segment, likely as a strategic decision to prevent overshooting column shifts along the homeward route.

### Underestimating Segments

No forager from Nest 1 or Nest 2 completed the full Leg 2 distance to the Elbow before turning (a few foragers from Nest 3 travelled the full distance to the 1^st^ Elbow), or the Leg 1 distance post-turn. These early turns are consistent with foragers either underestimating their vector segment distance or re-orienting before reaching the goal’s distance (the Elbow). Additionally, the orientation switch observed when full paths were analyzed (orientation from Leg 2 to Leg 1 rather than to the remaining global vector) suggests foragers were able to sequentially follow only the Leg 2 vector segment (pre-turn), while post-turn individuals only attended Leg 1’s vector segment of their global vector.

It is possible, though we believe unlikely, that this early turn away from Leg 2 is due to error accumulation within the path integrator system (leaky path integrator; See Collett and Collett 2009; Müller and Wehner 1988; Vickerstaff and Di Paolo 2005; Wehner 2020). Within the PI system, a portion of the accumulated distance estimate is gradually lost as distances increase, leading individuals to underestimate their homeward vector. However, this loss occurs as the PI system integrates paths over much larger distances than the 2.8m global vector these ants are accumulating. Additionally, in the current study, Nest 2 foragers (three-legged route) re-oriented back towards the 1^st^ Elbow compass direction after their second sharp turn, suggesting that they retained an estimate of that location even after their second turn, making the loss of the remaining Leg 2 vector unlikely. Finally, further evidence arguing against this hypothesis is observed in fan foragers, where they exhibited highly accurate fan vector paths to return to the column head (Freas et al. 2020). *V. pergandei* foragers following only their fan accumulated vector completed their full vector segment distance, beginning their search at the 1.5m vector distance. All evidence points to the potential for PI loss being negligible at these distances.

Instead, the pheromone’s absence during testing is likely a critical factor in the early turns away from the 1^st^ Elbow direction. In *V. Pergandei*, the pheromone’s presence confirms to foragers they are on the right route but have not overshot the nest, promoting continued vector orientation (Freas and Spetch 2021). Evidence that the pheromone trail acts as a reassurance cue during navigation has also been observed in other trail-following species (Wetterer et al. 1992; Czaczkes et al. 2011). The pheromone’s influence on the PI system is complex, playing a role in maintaining vector orientation, backtracking, and the vector segment attendance underlying the fan-and-column foraging structure (Freas et al. 2019ab, 2020; Freas and Spetch 2021). The pheromone’s presence influences the PI system to the extent that foragers which have no remaining vector, having run off their full global vector, still orient in the vector direction when on the pheromone (Freas and Spetch 2021), suggesting its presence is profoundly intertwined with how these animals navigate. *V. pergandei* foragers rarely orient to their column vector beyond ∼3 meters in the absence of this cue, even when tested after accumulating a column vector extending up to 24m (Plowes et al. 2019; Freas et al. 2019ab; 2020).

Given the pheromone’s importance to how these forager’s use their PI system across all these contexts, it appears likely it also underlies the decision to cut short the running of the vector segment. We hypothesize foragers’ early turns are a function of the pheromone’s absence during testing and this may act as an indicator that the forager has overshot the turn location. Foragers orient to their vector segment while a significant portion of it remains, even when not in contact with the pheromone as, in natural conditions, the column should continue this compass direction (Freas et al. 2020; Freas and Spetch 2021). However, as the vector segment is run down and uncertainty increases, the continued absence of the pheromone may be a signal that the forager has gone too far and should turn as most of the remaining column is in a new compass direction. This would make the behaviour very similar to backtracking in this species with foragers backtracking in the pheromone’s absence after completing over half their column vector, but well before reaching the end of their global vector (Freas et al. 2019a). This would also explain the differences we see in accurate distance estimation when testing fan foragers (Freas et al. 2020) compared to the current study, as fan foragers would always be traveling their full fan vector segment distance in the absence of the pheromone, thus its absence would not be an indication to search early.

### Limitations of vector segments

Our observed headings during three-legged route testing are less clear cut yet imply some interesting constraints. Initial headings for Nest 2, while oriented along Leg 3 and the global vector, could also be the result of orientation to the 1^st^ Elbow (combining Leg 3 and Leg 2). This leaves the possibility that attending a single non-nest waypoint within the column may represent a limiting factor within the column. This would be consistent with the lack of heading change observed between 30cm and the first sharp turn in foragers from Nest 2.

With Nest 3 foragers we do see significant heading changes between initial and first sharp turns, yet initial headings were only directed to Leg 3 in one of two conditions and could not be untangled from global-vector orientation in either, making such a distinction difficult. By the first turn in both conditions (*Column Head* and *Elbow Exposure*), foragers were well oriented to the compass direction of the 1^st^ Elbow (a combination of Leg 3 and Leg 2), once again showing evidence of a single waypoint within the column. Together, the lack of clear headings in the Leg 3 compass direction suggests foragers may struggle to orient along routes that contain multiple detours and the system which underlies attending a vector segment may only be able to waypoint foragers back to the 1^st^ Elbow rather than multiple turns along the route in sequence. The observed heading changes between Nest 3 and the lack of such a change with Nest 2 foragers (30cm vs. turn) may have been due to the larger directional shift preceding Leg 3 with the Nest 3 set-up (90° compared to 45° at Nest 2) or differences in the Leg 3 length (1m vs. 0.5m). These differences suggest the possibility that turn magnitude or segment length along the column plays some role in the separation of vector segments within the navigator’s memory or if one waypoint is a limitation of the system. Clearly, more work would be needed to pin down such mechanisms.

### Ecological Significance

Research on path integration in ants is primarily focused on solitarily foraging desert ants (*Cataglyphis* and *Melophorus*) which are diurnal, ‘thermophilic’ scavengers. Given this ecological niche, attending to a global PI to return along the shortest, straight-line distance to the nest, rather than retracing the outbound search, would likely be advantageous by minimizing the time spent exposed to the heat. When species instead forage as a group during periods where temperatures are moderate, maintaining the chemical connection to the nest via contact with the pheromone may be more advantageous, leading trail-following foragers to retrace the outbound route on their inward journey rather than leave the pheromone trail to travel along the global vector directly home. Evidence in *V. pergandei* suggests that, while the pheromone trail lacks polarity, it acts as a critical confirmation cue mediating the expression of multiple PI-based navigation behaviours (Freas et al. 2019a, 2020; Freas and Spetch 2021). Navigational behaviours often rely upon interactions between directional and non-directional cues. Given the heavy importance of the pheromone cue to this species’ PI system, this motivation to return to and maintain contact with the pheromone likely underlies the differences we observe in how *V. pergandei*’s PI system operates compared to solitarily foraging ants. For *V. pergandei,* leaving the pheromone is inhibited, meaning foragers should largely avoid global vector shortcuts if it results in leaving the pheromone. Instead, the PI system should direct movement in a way that keeps individuals in contact with the pheromone during the inbound route, leading the PI system to contain a mechanism to allow non-nest-based orientation to both re-enter the pheromone from the fan and to allow foragers to retrace curved outbound routes. At present, it remains unknown if this ability to retrace routes via the PI system alone exists more broadly across ants.

A final unanswered question that eludes our understanding in the current study, which concerns foragers’ post-turn paths during Nest 2 testing (2^nd^ path segment, Figure 4c) in the three-legged route. These foragers were highly directed yet not in any predicted Leg or vector direction, travelling perpendicular to the global vector and away from any point along the outbound route. We posit two possibilities. First, this may have been an attempt to orient to Leg 1 that was biased to the right of the true direction. However, we think this unconvincing as after this path segment, foragers still attempted to reach the 1st Elbow. Alternatively, foragers may have been attempting some form of search for the pheromone trail, which was abandoned after a short distance. This and many other questions merit further exploration.

### Conclusions

Inbound *V. pergandei* foragers navigate via distinct segments of their global PI to retrace separate legs of non-straight-line routes inconsistent with following a global vector. Attendance to vector segments was tested at an unfamiliar site, meaning the use of vector segments was unlikely to be externally triggered and more likely is integrated into their PI during outbound travel. We also show evidence that this ability may be limited within the pheromone column to one waypoint along the homeward route, with foragers along three-legged routes attending a combination of route segments. We propose that this way-pointing ability may set trail-following ant’s PI system apart from other species due to their foraging ecology, which encourages contact with the pheromone trail over direct nest-ward shortcuts.

## Declarations

### Ethics

There are no state or federal governmental regulations guiding the research of invertebrates in Arizona, USA. Manipulations were non-invasive and tested individuals returned to the nest to resume foraging.

### Availability of data and materials

The raw data for all testing conditions is publicly available online at OSF.IO.

